# *Ce*Aid: A smartphone application for logging and plotting *C. elegans* assays

**DOI:** 10.1101/2021.05.18.444737

**Authors:** Salman Sohrabi, Rebecca S. Moore, Coleen T. Murphy

## Abstract

*C. elegans* is used as a model organism to study a wide range of topics in molecular and cellular biology. Conventional *C. elegans* assays often require a large sample size with frequent manipulations, rendering them labor-intensive. Automated high-throughput workflows may not be always the best solution to reduce benchwork labor, as they may introduce more complexity. Thus, most assays are carried out manually, where logging and digitizing experimental data can be as time-consuming as picking and scoring worms. Here we report the development of *Ce*Aid, ***C. e**legans* **A**pplication for **i**nputting **d**ata, which significantly expedites the data entry process, utilizing swiping gestures and a voice recognition algorithm for logging data using a standard smartphone or Android device. This modular platform can also be adapted for a wide range of assays where recording data is laborious, even beyond worm research.

## Introduction

The nematode *C. elegans* is a powerful multicellular model organism that is used to study genetic pathways controlling cellular processes such as development, reproduction, and age-related decline. This is due to its small size, short generation time, genetic amenability, and conservation of cellular and molecular processes across species^1^. Many *C. elegans* protocols require a large sample size^2^. Much progress has been made in assay automation utilizing liquid workflows, robotic imaging platforms, and data analysis software. These tools enable users to carry out high-throughput genetic and chemical screens^3, 4^. For instance, advanced tracking algorithms coupled with imaging platforms were developed to automate data acquisition and analysis of *C. elegans* body shape and locomotion^5, 6^. WormFarm and WorMotel are also examples of microfluidic platforms that use automated algorithms for device operation, image processing, and data analysis focusing on expediting life span assays^7,8^. These tools are generally assay-specific, and the development of novel high-throughput techniques is frequently needed for new assays or protocols. Additionally, complete automation of some assays, such as fluorescence imaging and complex behavior tracking, may be much harder to achieve. In some cases, automation of experiments with recurring operational steps, such as assays of brood size and reproductive span, require daily data collection and cannot be easily scaled^9^. Microfluidics technologies have proven to be a powerful tool for handling complicated protocols. However, fabrication and operation of chips may require an engineering background and can be time-consuming, often halting the further use of these tools at the proof-of-concept technological validation step^10^. Overall, traditional methods for the culture of *C. elegans* on solid agar plates, picking worms using a platinum wire, and scoring them manually is an invaluable part of worm research. Thus, aiding researchers to expedite these manual tasks would be worthwhile.

In many standard manual assays with a large sample size, the researcher has to frequently switch between pick and pen to record data, searching for the proper location on the datasheet. The final step of digitizing a massive data log can often be as labor-intensive as performing the assay itself. To minimize the amount of time that the experimenter spends recording data, we have developed ***Ce*Aid** (***C**. **e**legans* **A**pplication for **i**nputting **d**ata) to automate the data entry process, which can be used for most manual assays in worms. Using features such as voice command, swiping, and tap gestures, *Ce*Aid does not require the user to shift focus to the pen and paper throughout the assay; all that is needed is a standard smart phone. We have utilized *Ce*Aid to log conventional *C. elegans* assays such as life span (LS), reproductive span (RS), brood size, and choice assays. The source code along with the APK file for *Ce*Aid are available on https://github.com/murphylab-Princeton/CeAid and https://murphylab.princeton.edu/CeAid.

## Results

Users can interact with *Ce*Aid through voice command, swiping, and tap gestures while the talk-back feature reconfirms the user’s input and declare the state of the app for receiving new data. Slider bars and embedded Numpad are also integrated into *Ce*Aid platform, allowing users to input data manually. New experiments can be set up by adding relevant information (**Figure 1a-b**). Scrolling through the expandable list of created experiments (**Figure 1c**) and tapping on any assay image, users are presented with a gallery view of color-coded icons showing the stored data or vacant slots (**Figure 1d-e**).

**Figure 1.**
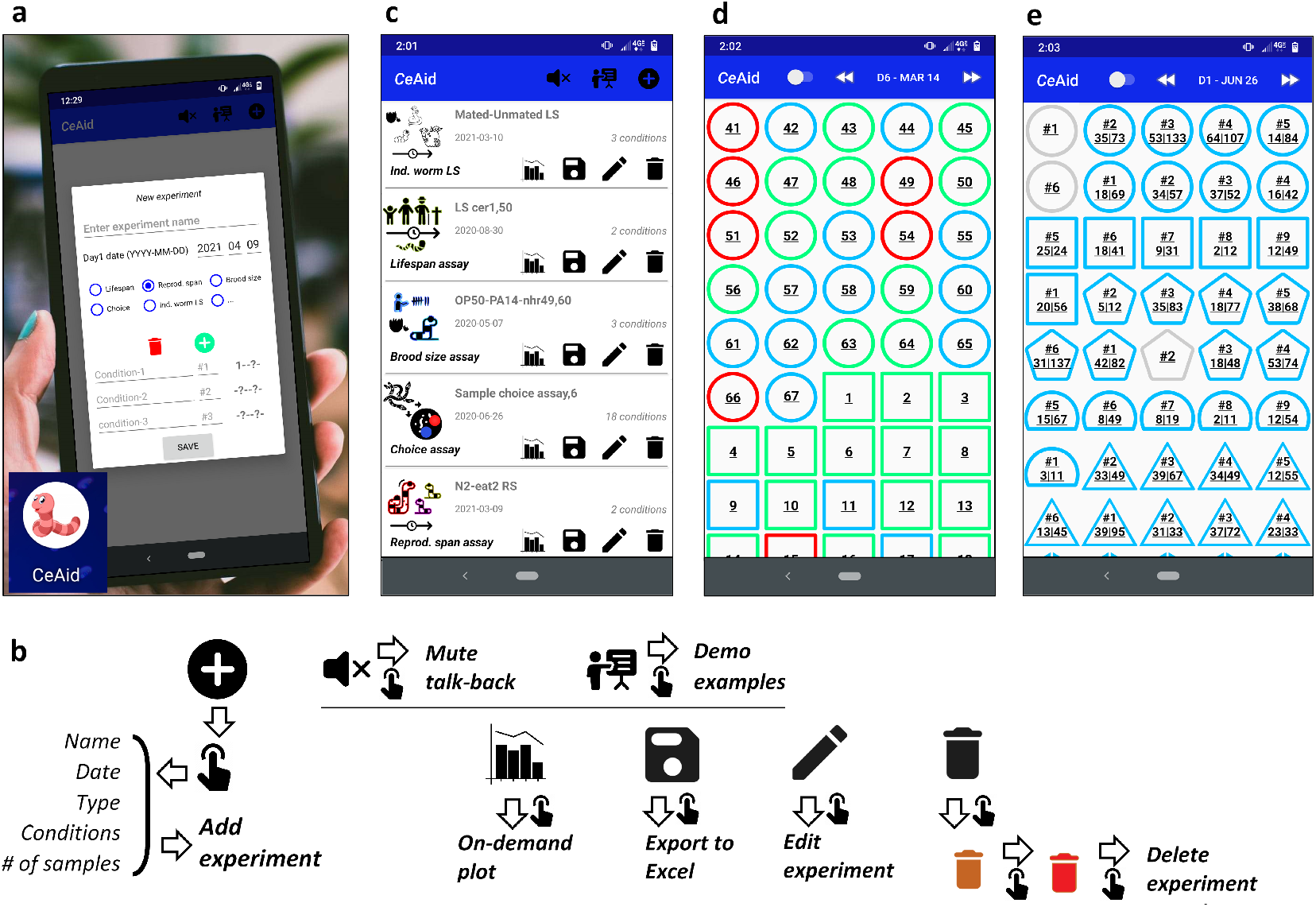
*Ce*Aid app. (a) New experiment pop-up. (b) The respective flow charts for creating and navigating experiments. (c) Expandable list-view of created experiments. (d) Color-coded gallery view of reproductive span assay. Toggle icon switch between numbering styles. Each condition is shown by a different shape. Green, blue, and red represents reproductive (RP), no progeny (NP), and censored (C) states of animals, respectively. (e) Gallery view of individual data points in the choice assay. Gray represents slots with no data. The values of left and right choices are shown on each item. 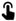 icon represents single-tap.

### Lifespan assay

*C. elegans* is a powerful model organism for studying aging due to its relatively short lifespan. To record data for LS assay, a voice recognition algorithm can be initiated with a double-tap anywhere on the screen to start listening for a two-digit number (**Figure 2a**). The talk-back feature confirms the user’s input after the sequential recording of the number of alive, dead, and censored animals. Horizontal and vertical swiping gestures navigate through samples and days of the experiment, respectively. Alternatively, users can input data manually using slider bars (**Figure 2b**).

**Figure 2.**
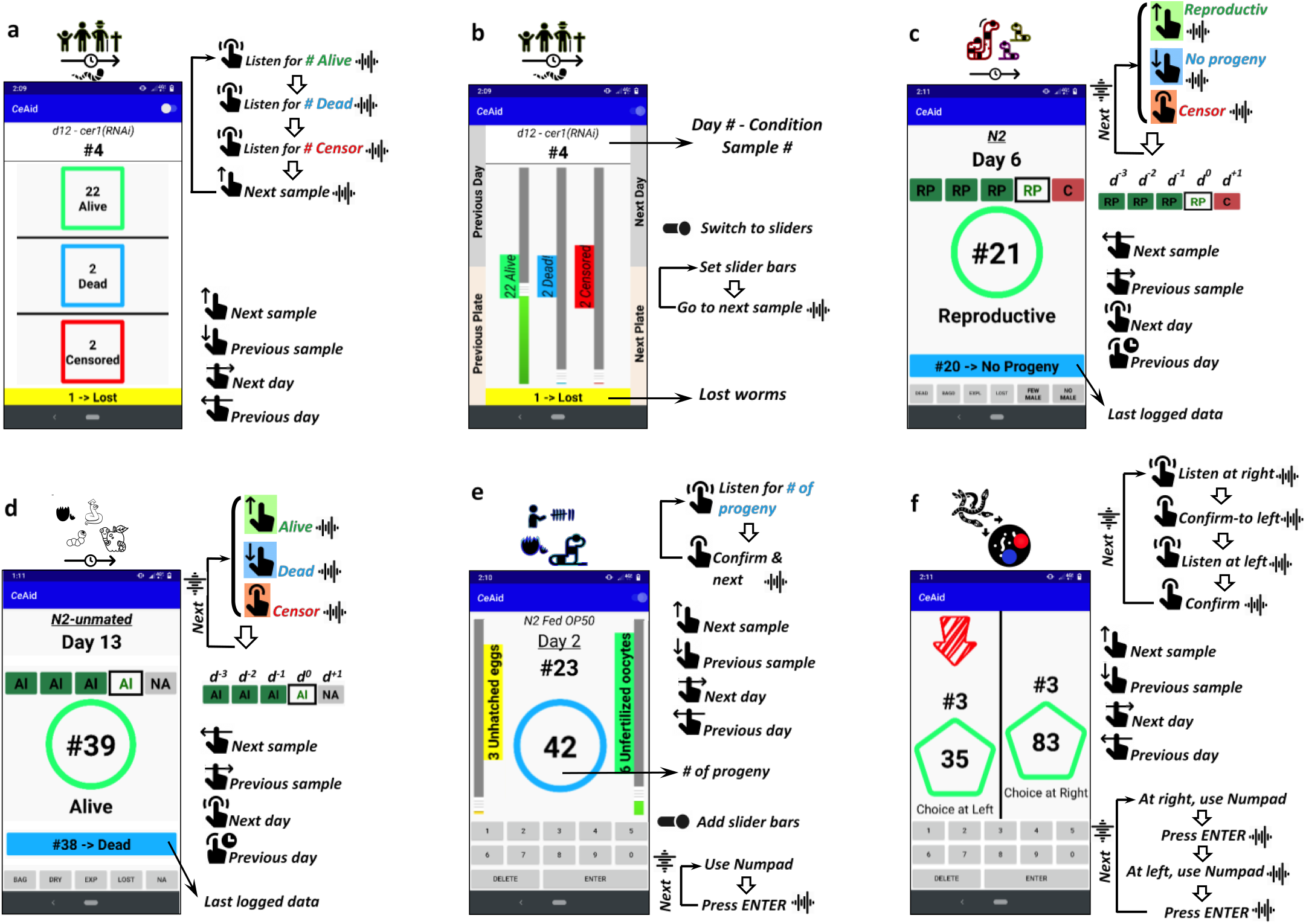
Data entry screens for various assays along with their respective flow charts demonstrating steps for logging data. Icons represent 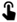 single tap, 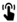 double tap, 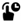 long press, 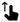 swipe up, 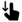 swipe down, 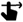 swipe right, 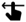 swipe left, and 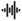 talk-back. (a) In automated lifespan assay, double-tap anywhere on the screen initiates the voice recognition algorithm to listen for the user’s input. (b) Slider bars in manual LS assay are also used to record # of alive, dead, and censored animals for each plate. (c) Users can record if an individual worm is reproductive on a specific day by swiping up and down. The most recent logged data, as well as the status of that animal in previous/next days, are also shown on the screen. (d) Similar to reproductive span assay, users can swipe up and down to report alive (Al) and dead (De) animals to input LS data for individual worms. The app skips dead and censored animals on subsequent days (e) Voice recognition program, Numpad, and slider bars can be utilized to record brood size assay data. (f) Choice assay data can be logged sequentially by activating voice recognition or using Numpad.

### Reproductive span assay

Female reproductive cessation is one of the earliest hallmarks of age-related decline. *C. elegans* has proven to be a useful model organism to better understand conserved mechanisms that regulate reproductive aging^11^. The duration of the progeny production period can be measured by checking whether hatched progeny exists on plates each day during the reproductive period. Swiping up or down on *Ce*Aid, the user can record if an individual worm is reproductive on a specific day, while a single tap can be used to censor the worm from the experiment, if needed (**Figure 2c**). More censoring options/notes are also provided at the bottom of the screen. After initiating these commands, the talk-back algorithm confirms input data and automatically prompts the user to the next sample. Swiping left and right enables navigation through samples and double-tap and long-press switch the day of the experiment. In LS assays where data are needed for individual worms, the app provides a similar platform as RS assay to instead log dead or alive states (**Figure 2d**).

### Brood size assay

The number of live progeny produced daily can also be used to study reproduction. Wild-type hermaphrodites use their limited number of self-sperm and can produce about 250 self-fertilized progeny, while mated animals can produce a substantially larger number^12^. Brood size assays within *Ce*Aid incorporate both voice recognition machine and Numpad to record the number of progeny produced (**Figure 2e**). Double-tap initiates listening algorithm while long-press censor the sample on a specific day. The number of unhatched eggs and unfertilized oocytes can also be recorded when slider bars appear on the sides of the screen. Similar to the LS assay, vertical and horizontal swiping provide navigation through samples and days of the experiment, respectively.

### Choice assay

In choice assays, animals choose between two spots on agar plates, which can be used to measure behavioral phenotypes such as dietary preference^13^, chemotaxis^14^, olfactory learning behavior^15^, and avoidance of pathogenic bacteria^16^. To record choice assay data, the user can activate the voice recognition algorithm by double-tapping on screen or, alternatively, use an embedded Numpad at the bottom of the screen (**Figure 2f**). Furthermore, swiping gestures enable navigation through recorded samples.

### Visualization

Recorded data can be plotted within *Ce*Aid or exported to an Excel file for further statistical analysis (**Figure 3**). Using on-demand plotting, the trajectory of the experiment can be visualized while it is happening prior to analysis using statistics programs (**Figure 3a**). The demo examples are also integrated into the app and can be produced on the main activity page.

**Figure 3.**
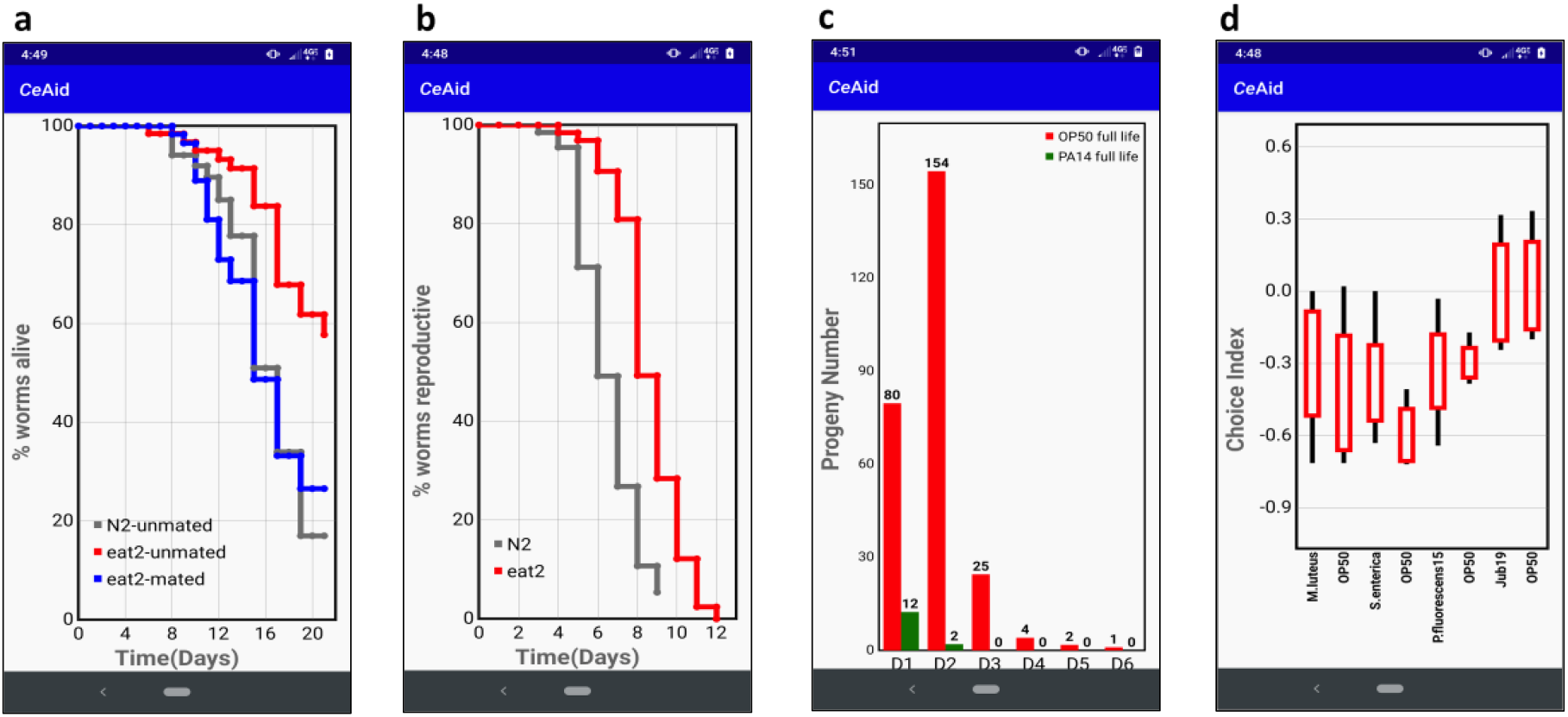
On-demand plotting. (a) Survival curve of an undergoing experiment for sample populations of mated and unmated animals. (b) The reproductive span of self-fertilized N2 and *eat-2* worm strains. (c) Wild-type worms exposed to P. aeruginosa (PA14) beginning on Day 1 of adulthood produce less progeny compared to whole-life OP50 fed worms. Swiping left/right navigates to plots for daily number of unhatched eggs and unfertilized oocytes. (d) Wild-type worms were trained on different pathogenic bacteria. Following training, an aversive learning assay was conducted to determine worms’ food preference compared to control food^17^. Choice index = (number of worms on OP50 – number of worms on bacteria)/(total number of worms). The box plot shows minimum, 25th percentile, median, 75th percentile, maximum. The number of conditions to be plotted can be set and swiping left/right provides the capability to navigate through them.

## Conclusion

*Ce*Aid is developed to automate manual data entry procedures for *C. elegans* assays through a series of highly simplified steps. This opensource android application is modular and functions for tap and swipe gestures along with voice recognition algorithm, slider bars, and embedded Numpad can be adapted for newer scoring assays. Overall, eliminating the need to shift the user’s focus to pen and paper throughout the experiment can significantly expedite worm research.

## Acknowledgments

S. S. and C.T.M. designed the layout for the app. S. S. developed the code. R.S.M. provided demo data. S. S. and C.T.M. wrote the manuscript. C.T.M. supervised the project and acquired funding. We thank Murphy lab for discussion, particularly Rachel Kaletsky, Cheng Shi, Vanessa Cota, Will Keyes for great suggestions and sharing data. This work was supported by the Glenn Foundation for Medical Research award to C.T.M. (GMFR CNV1001899) and NIH award RF1AG057341 (NIA).

